# Decreased snow cover stimulates under ice primary producers, but impairs methanotrophic capacity

**DOI:** 10.1101/380170

**Authors:** Sarahi L Garcia, Anna J. Szekely, Christoffer Bergvall, Martha Schattenhofer, Sari Peura

## Abstract

Climate change scenarios anticipate decrease of spring snow cover in boreal and subarctic regions. Forest lakes are abundant in these regions and substantial contributors of methane emissions. We performed an experiment on an anoxic frozen lake and observed that the removal of snow increased light penetration through the ice into the water modifying the microbial composition across depths. A shift in photosynthetic primary production was reflected by the increase of chlorophyll a and b concentrations in the upper depths of the water column, while Chlorobia, one of the key photosynthetic bacteria in anoxic lakes, shifted to lower depths. Moreover, a decrease in abundance of methanotrophs, such as Methylococcaceae, was noted concurrently to an increase in methane concentration in the water column. These results indicate that decrease of snow cover impacts both primary production and methane production/consumption, ultimately leading to increased methane emissions after spring ice off.

## Introduction

Small forest lakes are a typical feature of the boreal and subarctic region ^1^. These small water bodies with high organic loads are hotspots in the carbon cycle and one of the most prominent environmental sources of greenhouse gas emissions in these regions ^2,3^. The microorganisms inhabiting such lakes are the main drivers of these biogeochemical processes ^4^. However, the community structure and functioning of microorganisms is bound to the environmental conditions and seasonality. One very important factor related to seasonality is light availability. Light is the driving force of primary production at the base of the food web. Algae provide substrates and oxygen for lake bacteria and are an important food source of zooplankton. In wintertime, ice cover inhibits oxygen transfer from the atmosphere into the water column while snow cover impedes light transfer, further curtailing photosynthesis beneath the ice ^5,6^. Moreover, aerobic microorganisms consume residual oxygen in the water beneath the ice ^7^, leading to decreasing oxygen gradient and anoxia from the lake surface to the bottom. The resulting anoxic conditions facilitate anaerobic processes like methanogenesis and decrease methanotrophy ^8^ These conditions result in methane accumulation under ice and consequently high methane emissions during ice break-up in the spring ^9,10^. Currently, climate change is altering seasonal patterns in the subarctic region ^11^ and also changing patterns of snow cover worldwide ^12^. In the northern hemisphere these changes include sudden extreme conditions, as seen during the 2007/08 winter, when a sudden loss and reformation of snow cover occurred due to a warming event ^13^. Also long term trends have been observed, such as a decrease in spring snow cover ^12^. This project tests the impact of reduced snow cover on a humic lake ecosystem with the emphasis on methane cycle and primary producers.

We hypothesized that (a) the decrease in snow cover increases light penetration and stimulates primary production. Furthermore, we hypothesized that more primary production would (b) increase the availability of oxygen having an impact on aerobic bacteria and (c) increase methanotrophic activity as an effect of improved oxygen availability. The project was conducted as a whole lake manipulation in Lake Lomtjärnen (63°20′56.9″N 14°27′28.3″E) a small boreal lake in central Sweden.

## Results and Discussions

### Snow cover removal increased light penetration and its effect on chlorophyll

Our sampling scheme included 6 depths (depth 1 was sampled at 0.65 cm and thereafter approx. every 40 cm) and 6 time points. We sampled a vertical profile of the lake on three occasions before the snow removal and three times after; with one day between each sampling. Thus, the total duration of the experiment was two weeks. Snow depth on the frozen lake was 18 to 21 cm through the experiment. The snow was removed from approximately an area of 400 m^2^ in the middle of the lake. During the experiment the ice thickness was approximately 50 cm. Prior to the snow removal, the lake had a shallow epilimnion, with a steep oxygen depletion layer from 0.6 to 0.7 m, where the oxygen concentration was close to detection limit (Figure S1).

We observed light intensity increase in all depths of the lake after the snow removal (depth 1 from 177 to 3764 lx, depth 2 from 63 to 1378 lux, depth 3 from 39 to 826 lux, depth 4 from 20 to 480 lux, depth 5 from 9 to 299 lux and depth 6 from 0 to 112 lux). Light intensity was significantly higher only in the first three layers (0.65 m to 1.35 m) (Figure 1). Moreover, the chlorophyll a concentration increased significantly in those top three layers (0.65 m to 1.35 m), whereas chlorophyll b increased in layers 4 and 5 (1.85 m to 2.53 m). In comparison, the concentration of bacteriochlorophyll d and e appeared to increase in depth 5 (Figure 1). Based on these results we presume light had a direct effect on the phytoplankton of the water column affecting their chlorophyll production and photosynthesis activity and subsequent oxygen production. After the snow removal an increase in oxygen concentration in the top layer under the ice was observed which rapidly decreased as time progressed (Figure S1). However, the differences in oxygen concentration were not statistically significant. This might indicate that heterotrophic organisms immediately utilized the produced oxygen, which is supported by the fact that bacterial abundance doubled in the upper layer of the lake (Figure 1).

**Figure 1.**
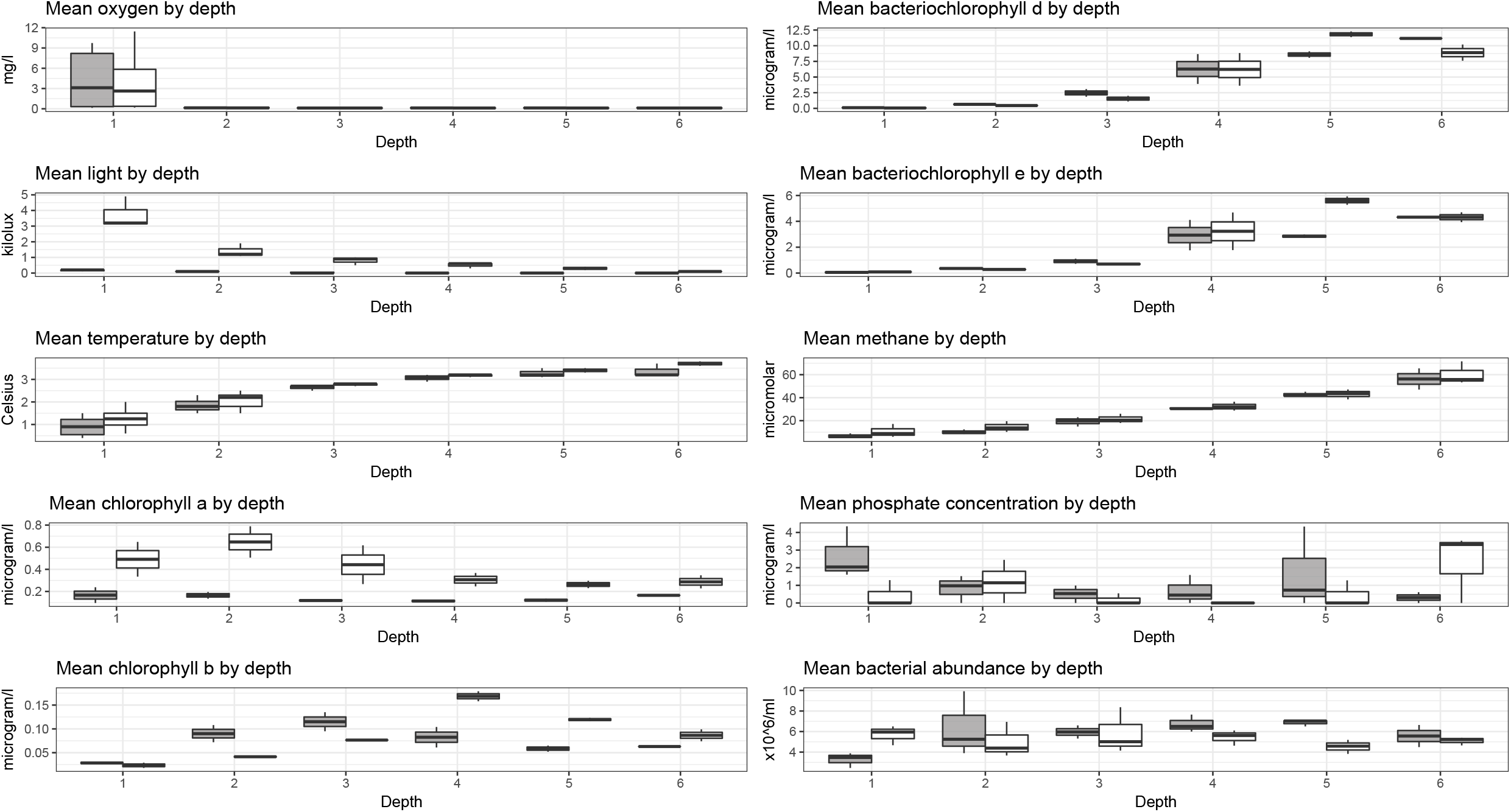
Chemical and physical parameters of the lake before (grey) and after snow removal (white). Depth 1 = 0.65 m, 2 = 1.0 m, 3 = 1.35 m, 4 = 1.85 m, 5 = 2.4 m, 6 = 2.7 m

### Microbial community changes

Against our expectation, snow removal did not have a significant effect on the total community composition (PERMANOVA: p_DNA_ = 0.594, p_RNA_ = 0.636). However, when the depths were considered individually, the composition of microbial communities before and after snow removal were significantly different in depth 1 (0.65 m) and in depth 4 (1.85 m) (PERMANOVA p_depth1_ = 0.007 and p_depth4_ = 0.046). Depth was an important driver of community composition (PERMANOVA, p < 0.001), as oxygen concentration and redox potential are key factors structuring bacterial communities ^14,15^. The major microbial groups in the first depth were Betaproteobacteria (26% before and 32% after snow removal in relative abundance of DNA-based bacterial community composition, 30% before and 46% after snow removal in RNA-based community) and Gammaproteobacteria (23% before and 7% after snow removal in DNA, 37% before and 18% after snow removal in RNA) (Figure 2). Most of the betaproteobacterial OTUs were classified as Comamonadaceae, while most of the gammaproteobacterial were classified as Methylococcaceae. Parcubacteria also contributed to the community change post snow removal in depth 1 (5% before and 10% after snow removal in DNA, not present in RNA). The major microbial groups in the fourth depth were Chlorobia (33% before and 17% after snow removal in DNA, 40% before and 28% after snow removal in RNA), Chloroflexia (12% before and 10% after snow removal in DNA, 13% both before and after snow removal in RNA) and Anaerolineae (3% before and 4% after snow removal in both DNA and RNA) (Figure 2). Changes in both DNA and RNA were consequent in most of the groups in which we observed the biggest alterations post treatment. Overall, the 11 most abundant OTUs contained each at least 1% of the sequences. Together these 11 OTUs accounted for 32% of the sequences. The most abundant OTU belonged to the Chlorobiaceae family, while the second and third most abundant OTUs belonged to the Methylococcaceae family.

**Figure 2.**
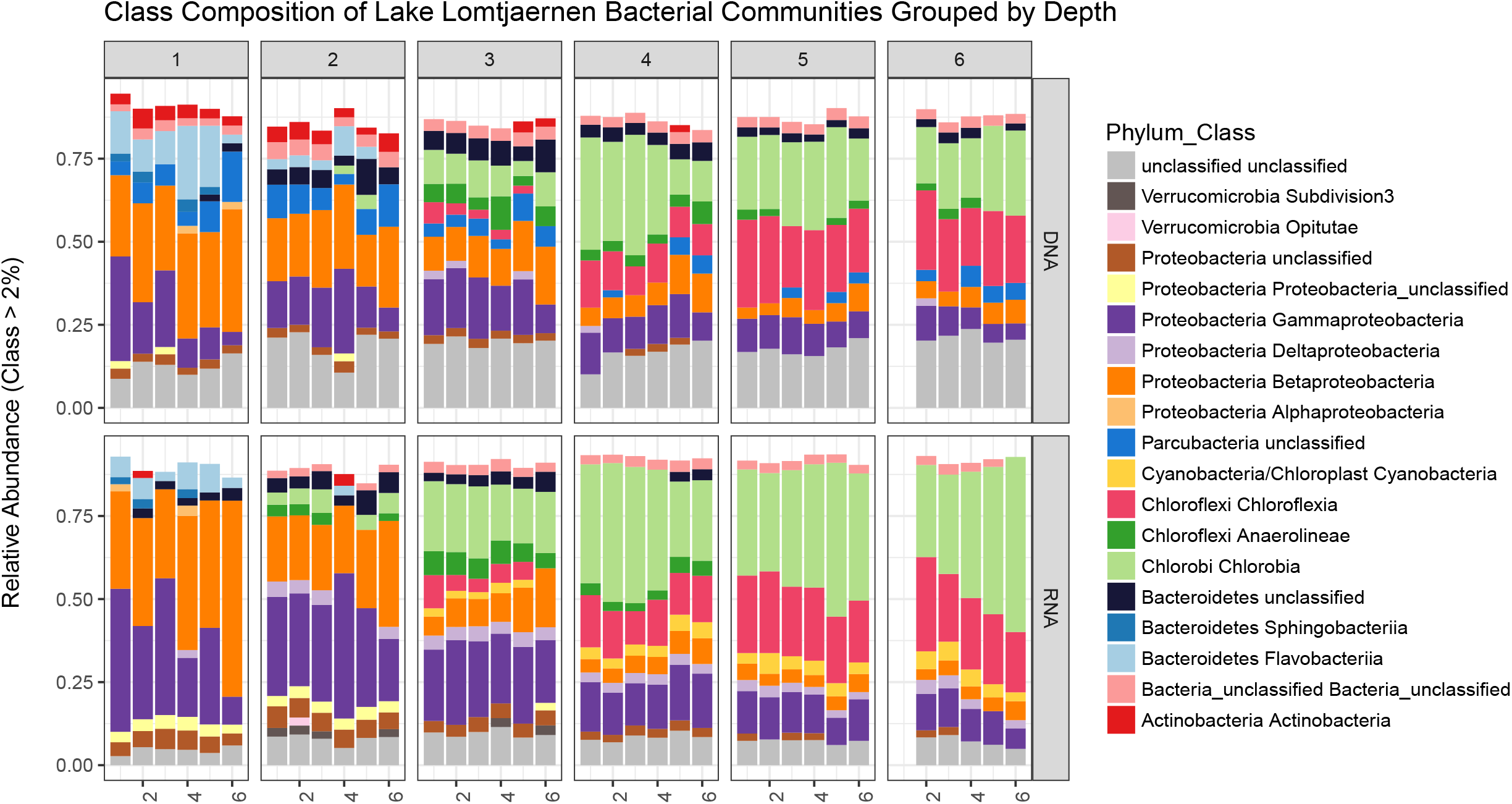
Relative abundance of microbial communities of each of the samples. Color coded according to taxonomical class. In the x axis 1 = 100%. Numbers in upper boxes represent the depth number (1 = 0.65 m, 2 = 1.0 m, 3 = 1.35 m, 4 = 1.85 m, 5 = 2.4 m, 6 = 2.7 m). Bars inside each box are ordered according to time being the first bar the time 1 and the last bar time 6. First bar in depth 6 is missing as on the first day of sampling we could not obtain a sample for depth 6.

Because we expected the oxygen conditions to improve the fastest in the top layer of the water column, we looked into the most abundant families in the first depth. We observed an increased relative abundance of the family Comamonadaceae (15% to 19% in DNA, 18% to 30% in RNA) and Flavobacteriaceae (10% to 14% in DNA, 5% to 6% in RNA) (Figure 3). Comamonadaceae and Flavobacteriaceae are both aerobic heterothrophs ^16-18^. Both of these organisms that increased in relative abundance after snow removal could take advantage of the increased availability of oxygen and organic compounds originating from increased activity of the primary producers, while keeping the oxygen concentration seemingly unchanged.

**Figure 3.**
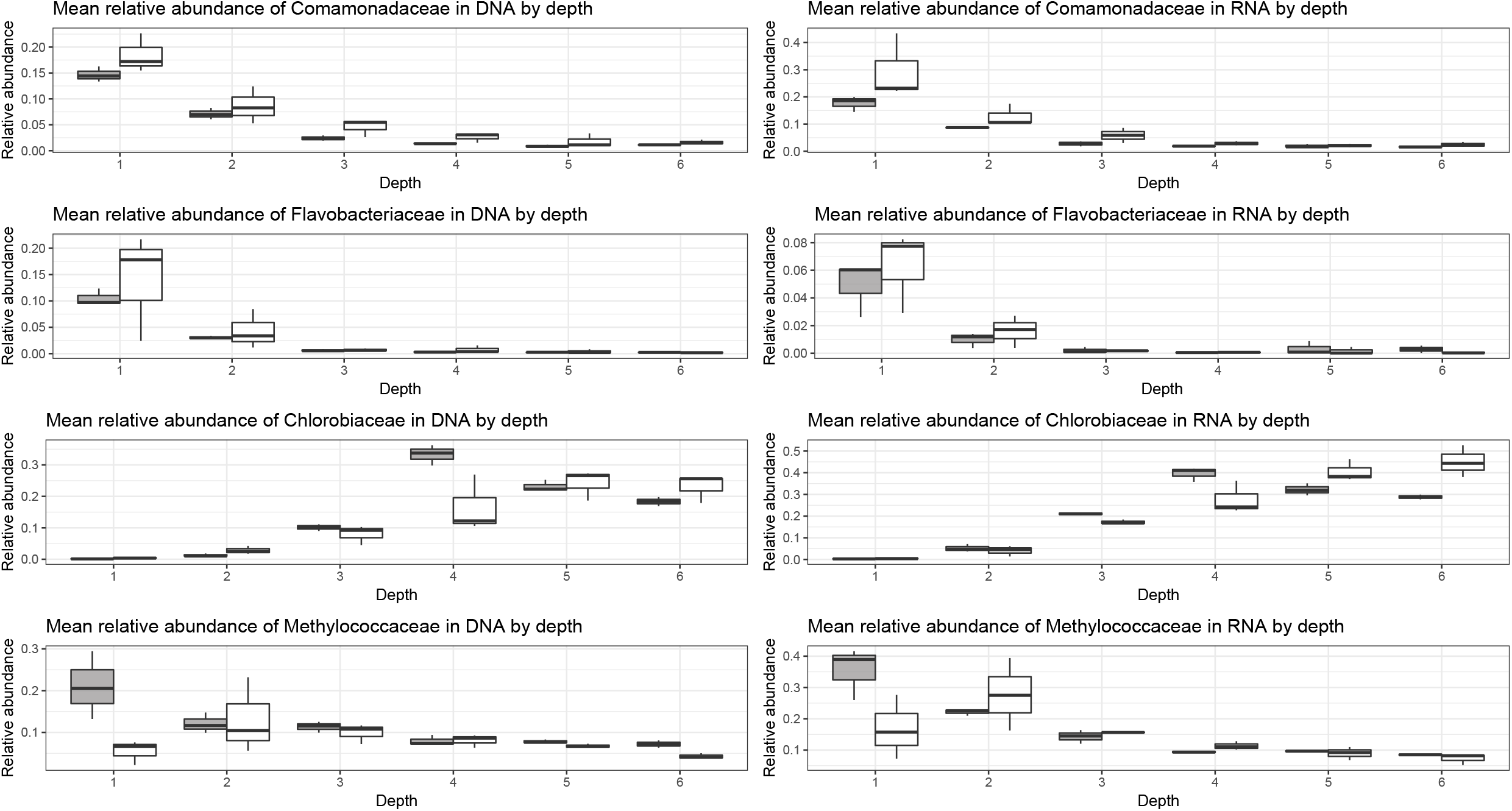
Relative abundance of some specific taxonomical groups. Samples taken before the snow removal are in grey and after snow removal in white. In the x axis 1 = 100%. Numbers in depth number; 1 = 0.65 m, 2 = 1.0 m, 3 = 1.35 m, 4 = 1.85 m, 5 = 2.4 m, 6 = 2.7 m.

Chlorobia are photoautotrophic bacteria that are common in boreal lakes such as our study lake ^19-21^. Because of the extraordinarily large numbers of bacteriochlorophylls in their antenna complexes, some of these green sulfur bacteria are able to grow at extremely low light irradiances (1–10 nmol photons m^−2^ s^−1^) under which no other types of chlorophototrophs can grow ^22,23^. It is possible that the decrease in the abundance of Chlorobia in depth 4 after snow removal was due to light intensity increase, making the conditions more favorable for organisms using chlorophyll a and b (Figure 1). Moreover, an increase of Chlorobi both in DNA and RNA relative abundances was observed in depth 5 and 6 of the lake (Figure 3). This together with the increase in bacteriochlorophyll d and e in depth 5 could mean that decreased snow cover switches the taxonomical composition of the primary producers of the lake pushing the optimal conditions for Chlorobia to lower depths of the lake. This shift in primary producers might have implications for the carbon balance of the lake, following the possible shifts in efficiency of carbon dioxide uptake and microbial interactions.

### Methane after snow removal

As expected methane concentration increased with depth, but opposite to our hypothesis methane concentration was significantly higher after the snow removal (two-way ANOVA p_depth_ < 0.001, p_treatment_ = 0.020) (Figure 1). Most of the gammaproteobacterial organisms observed in the water column belonged to the family Methylococcaceae, which consists solely of methane consuming bacteria. Methylococcaceae decreased in relative abundance in DNA after the treatment on all of the depths except for depth 4 (Figure 3). We also observed a decreasing trend in phosphate concentration in depths 1, 3, 4 and 5 after the treatment. In a previous study, low phosphate concentration has been linked to impaired methanotrophic activity ^24^. Considering that many methanotrophs are slow growing ^25^ and the increased algal activity in the lake after snow removal, as suggested by the increased chlorophyll concentrations, it is possible that the methanotrophs were unable to compete for phosphorus with the phytoplankton. Alternatively, the increased primary production could have contributed more substrate to methanogenesis in surface sediments, increasing overall the concentration of methane in the water column ^10^. Decreased methanotrophy and increased methanogenesis would explain the increased methane concentrations after the snow removal. In any case, increased methane concentrations in the water column translate into increased methane emissions to the atmosphere once the ice melts in the spring.

### Conclusions

Our results show that the impact of decreased snow cover to the lake microbiome and also to total lake metabolism is a complex interaction between the biotic and abiotic lake characteristics. While our first hypothesis (a), the increase of light and stimulation of primary production following the snow removal proved to be correct, (b) the increase of oxygen and (c) increase of methanotrophic activity appeared to be false. However, it should be noted that the duration of our experiment was rather short and it might be possible that on a longer time scale the oxygen conditions might improve. Nevertheless, we conclude that the final outcome of the decreased snow cover can be expected to change the total lake metabolism and potentially increase methane emission after the ice break.

## Methods

The experiment was conducted in March 2016 on Lake Lomtjärnan (63°20′56.9″N 14°27′28.3″E), a small lake located in Krokom (Sweden). The lake is located on a mire surrounded by a coniferous forest and it is ice covered during the winter months, approximately from November to April. The experiment consisted of two parts; monitoring of the lake while there was a snow cover over the whole ice and observing the impact snow cover removal after the snow was removed from an area of approximately 400 m^2^ in the middle of the lake. Snow removal was performed on day 6 manually using snow shovels. For each sampling a hole was drilled through the ice on a different location to avoid interference of unnatural ice formation as the holes were always filled and covered with snow to limit oxygen and light penetration to the water column.

Light intensity was measured with 18 HOBO loggers with the capacity to measure light intensity and temperature (HOBO Pendant^®^ Temperature/Light 64K Data Logger, Onset, USA). The loggers were placed under ice from surface to the bottom every 0.1 to 0.75 m on the first day of the experiment and kept there throughout the experiment. Laterally, the sensors were approximately 50 cm from the hole in the ice. The light values are presented as daily averages for the time between sunrise and sunset (approximately 10 AM to 3 PM).

On each sampling occasion samples were taken for the measurement of chemical parameters (Chlorophylls, Nitrite, Nitrate, Phosphate, Sulfate, Ammonia, Fluoride, Chloride, O2, CH4, CO2), and DNA- and RNA-based community analyses. For DNA and RNA the samples were taken with Sterivex-filters (Millipore, Billerica, MA, USA). For RNA the filtering was limited to 15 minutes, while for DNA the filtering was continued until the filter was clogging. Samples were taken from six different depths and on days 1, 3 and 5 (before the snow removal) and on days 7, 9 and 12 (after the snow removal).

Chlorophyll pigments were extracted on ice in 2ml 90% acetone using an ultrasonic bath and left at −20°C overnight. After that, the samples were centrifuged (3000x 10min at 4°C) and filtrated (0.45μm syringe filters). The extracts were analyzed using HPLC on an Agilent 1100 Series HPLC system (Agilent Technologies, Waldbronn, Germany) fitted with three RP-18e 100×4.6mm Chromolith Performance columns (Merck, Darmstadt, Germany) connected in series. The flow rate was 1.4ml/min and the gradient program followed as described ^26^. The column temperature was 25°C and the injection volume was 100μl (70μl sample + 30μl 0.5M ammonium acetate). The absorbance was measured with a diode array detector between 300-800nm (resolution 2nm and slit with 4nm). Chlorophyll a and b were identified and quantified using standard solutions (DHI Laboratory Products, Hoersholm, Denmark) and the bacteriochlorophylls by using previously published chromatograms, spectra and extinction coefficients ^27-31^. Methane concentration was analysed as described, except that room air was used instead of nitrogen for the headspace. The methane concentration was analysed also from room air and subtracted from the final gas concentrations.

DNA and RNA was co-extracted using phenol-chloroform method ^33^ as modified in ^21^. RNA was then transcribed into cDNA as previously descriped ^34^ using RevertAid H Minus First Strand cDNA synthesis kit (Thermo Scientific). After this, both, the RNA and DNA samples were amplified for bacterial 16S rRNA genes using primers 341r and 805f ^35^. PCR protocol was done as previously ^36^. The samples were then pooled in equimolar amounts and sequenced with Illumina MiSeq at Science for Life Laboratories (Uppsala, Sweden). The resulting 2.5 million sequences were processed using Mothur ^37^ as described ^38^, except that OTU clustering was done using abundance-based greedy clustering. Raw sequences have been submitted to ENA under the accession number ERS2597919 - ERS2597988.

The effect of snow removal and water depth was tested by two-way analyses of variance (ANOVA) with corresponding Tukey’s post-hoc tests for the environmental parameters and by two-way permutational ANOVA (PERMANOVA) with 9999 permutations for the community composition data. The normal distribution of the residuals of the ANOVA models was tested using Shapiro-Wilk normality test and where needed, data was log transformed. All statistical analyses were done in R version 3.4.3 ^39^. Packages phyloseq, vegan and ggplot2 were used ^40-42^.

## Author Contributions

SLG and SP originated the research. SLG, AS, MS, SP participated in the sampling campaign. SP, AS and CB conducted the laboratory work including DNA, RNA, nutrient, and chlorophyll analyses. SLG, AS, SP performed the data analysis. All authors contributed to discussions of data and manuscript review.

## Acknowledgements

This work was supported by grants from Olsson-Borgh foundation (grant to SP and SLG), the Academy of Finland (grant 265902 to SP) and the Royal Swedish Academy of Sciences (grant BS2017-0044). The computations were performed on resources provided by SNIC through Uppsala Multidisciplinary Center for Advanced Computational Science (UPPMAX) under Project SNIC 2017/1-616. The authors thank the shoveling team: mainly Zoltan Török, Philipp Baur and Jussi Peura.

---

**Supplementary Figure S1.** Oxygen concentration in the water column across the experiment in the sampling depths.

